# Coupling microalgal bioremediation of recirculating aquaculture effluents with photobiological hydrogen production

**DOI:** 10.64898/2026.01.15.699656

**Authors:** Marco L. Calderini, Valéria Nagy, Szilvia Z. Tóth, László Kovács, Soujanya Kuntam, Pauliina Salmi

## Abstract

Recirculating aquaculture system (RAS) effluents contain substantial nitrate and phosphate loads, posing a significant eutrophication risk to aquatic ecosystems if left untreated. While microalgal bioremediation is a promising strategy, expanding opportunities for valorisation is crucial for its successful implementation. This study couples the cultivation of nitrate-utilizing *Chlamydomonas reinhardtii* strain CC-1690 in RAS effluent with photobiological hydrogen (H_2_) production to achieve nutrient removal, bioenergy production, and high-quality biomass. Within 70 h, CC-1690 achieved near-complete depletion of nitrate and phosphate from the effluent. Subsequently, H_2_ production was induced via carbon limitation and anoxia, yielding a cumulative 49 µmol H_2_ mg^−1^ Chl over 72 h. Although decreased chlorophyll content and F_v_/F_m_ indicated physiological stress, key photosynthetic subunits (PsbA, PSBO, CP47, PetB and PsaA) remained largely stable. Importantly, the process preserved biomass quality; total lipid content increased slightly, enriched in palmitic (16:0) and α-linolenic (18:3ω-3) fatty acids while protein content was unaffected. These results demonstrate that integrating H_2_ production with RAS effluent remediation offers a robust circular economy approach, valorising wasted nutrients into bioenergy and high-quality biomass with potential downstream applications.

## 1. Introduction

Over the past decades, global aquaculture production has undergone rapid growth as a means of satisfying rising protein demand (FAO 2024). Recirculating aquaculture systems (RAS) have emerged as a sustainable approach to intensive aquaculture, in which closed-loop systems facilitate water use efficiency, improve biosecurity, and enable the optimization of production conditions (Bregnballe, 2022). Although water recirculation rates can reach 90–99% (Ahmed & Turchini, 2021), wastewater generation is unavoidable, and the accumulated volume becomes substantial in medium-to large-scale operations. This effluent contains high levels of nitrate and phosphate, primarily derived from metabolic waste and the dissolved portion of uneaten feed (Wang *et al*., 2012). Currently, nutrient-rich effluents from RAS are often released into the environment, presenting a significant eutrophication risk to aquatic ecosystems. While typically viewed as waste, these nutrients represent valuable resources. For instance, nitrogen fertilizers are energy-intensive to produce, contributing to greenhouse gas emissions, while phosphorus is sourced from finite phosphate rock reserves that are being rapidly depleted (Cordell *et al*., 2009). Recovering these nutrients from RAS effluents, therefore, offers a dual benefit: mitigating environmental pollution and recycling critical nutrients for further utilization.

Among various wastewater treatment strategies, microalgal bioremediation is a promising, eco-friendly solution for nutrient recovery and valorisation (Calderini *et al*., 2021; Böpple *et al*., 2024; End *et al*., 2024). Most microalgae are capable of assimilating nitrate and other inorganic nutrients present in RAS effluents into biomass through photosynthetic growth. Compared to established technologies for the treatment of RAS effluents, such as anaerobic digestion and pyrolysis, microalgal bioremediation does not require dewatering of the effluent (Brod *et al*., 2017), thereby reducing energy input and increasing overall process sustainability. Due to its high nutrient content and ratios compatible with microalgal requirements, as well as the absence of toxic chemicals, RAS effluent has been shown to outperform other waste streams, such as municipal or industrial wastewater, when used as a growth medium (Silkina *et al*., 2019). Microalgal production utilizing aquaculture effluents represents a multifunctional opportunity with diverse potential applications in the bioeconomy (End *et al*., 2024). Untreated biomass can be used as a biofertilizer (Coppens *et al*., 2016), while high-value pigments, proteins, fatty acids, polysaccharides, and bioactive metabolites can be extracted for use in a wide range of bio-based products, including plant biostimulants and aquafeed additives (Viegas *et al*., 2021; Shastak & Pelletier, 2023). Biomass quality is therefore key to enabling the valorisation of the recovered nutrients from RAS effluents.

Photobiological hydrogen (H_2_) production represents a promising strategy to expand the biotechnological application of photosynthetic microorganisms, particularly microalgae. H_2_ is a versatile energy carrier with a wide variety of industrial applications. Certain green microalgae species, such as *Chlamydomonas reinhardtii,* are capable of producing H_2_ as a byproduct of the photosynthetic electron transport chain, a process catalyzed by [FeFe]-hydrogenase enzymes (HydA1 and HydA2) located at the acceptor side of photosystem I (PSI) (Forestier *et al*., 2003; Sawyer & Winkler, 2017; Engelbrecht *et al*., 2020). These hydrogenases act as electron sinks, alleviating excess reductant accumulation and preventing over-reduction of the photosynthetic machinery during dark-to-light transitions (Godaux *et al*., 2015). However, once the Calvin-Benson cycle is operative under light, electrons derived from water splitting in photosystem II (PSII) are primarily utilized for carbon fixation (Godaux *et al*. 2015), while oxygen (O_2_) buildup irreversibly inactivates the hydrogenases (Swanson *et al*., 2015).

Strategies such as nutrient deprivation (typically sulphur) (Melis *et al*., 2000), balancing cellular respiration and O_2_ production (Elman & Yacoby, 2022), and establishing algae-bacteria co-cultures (Lakatos *et al*., 2014) maintain low O_2_ concentrations under photoheterotrophic conditions. Meanwhile, pulse illumination reduces the activity of the Calvin–Benson cycle under both photoautotrophic and photoheterotrophic conditions (Kosourov *et al*., 2018). Direct removal of O_2_ via scavengers coupled with substrate limitation of the Calvin–Benson cycle (CO_2_- and acetate-free conditions) is another strategy for sustained H_2_ production under photoautotrophic conditions (Nagy *et al*., 2018). This approach minimizes both O_2_ accumulation and the activity of the Calvin–Benson cycle, offers a high theoretical efficiency of converting light into chemical energy (11-13%), is applicable at sunlight intensities (Nagy *et al.,* 2021), and can avoid the degradation of photosynthetic complexes or cell death associated with nutrient-deprivation strategies.

In addition to the competition of the Calvin-Benson cycle with H_2_ production, nitrate assimilation acts as a potent alternative electron sink, specifically during the reduction of nitrate and nitrite, thereby suppressing H_2_ evolution (Barea & Cardenas, 1975; Aparicio *et al*., 1985). Given that only a subset of *C. reinhardtii* strains are capable of assimilating nitrate (Pröschold et al., 2005) and that H_2_ production studies are typically carried out in ammonium-containing media, the feasibility of coupling nitrate-containing media with H_2_ production has remained largely unexplored.

In this study, we demonstrate that the nitrate-utilizing *C. reinhardtii* strain CC-1690 efficiently removes nitrate and phosphate present in RAS effluents, unlike standard H_2_-producing strains (CC-124, T222^+^, *pgr5*) which lack this capability. Following nutrient removal, H_2_ production was induced directly in the effluent using the anaerobiosis-induced photoautotrophic protocol (Nagy *et al*., 2018). Although H_2_ yields were slightly lower compared to those in standard algal media, substantial production was obtained. Furthermore, despite the observed stress responses, the biomass quality was preserved, supporting downstream valorisation. These findings demonstrate the feasibility of integrating microalgal bioremediation of RAS effluents with photobiological H_2_ production using a nitrate-assimilating *C. reinhardtii* strain.

## 2. Materials and methods

### 2.1. Recirculating aquaculture system effluent

RAS effluent samples were obtained from the Natural Resources Institute Finland (LUKE) Laukaa fish farm (Pulkkinen *et al*., 2021), Finland. Effluent samples were collected during May of 2025 from the water outlet after drum filtration and fixed bed bioreactor treatment. The fish farm sources its water from an oligotrophic lake (lake Peurunka; 62.44886, 25.85201), and the sampled effluents originate from a RAS system which at the time of sampling was used for low-density rearing of rainbow trout (*Oncorhynchus mykiss*). After collection, effluent was filtered using tangential-flow filtration (Millipore Durapore cassette, 0.22 µm pore size), UV-sterilized for 45 min and stored in the dark at 4 °C.

### 2.2. Algal growth conditions and H_2_ production

*C. reinhardtii* wild-type strains CC-1690 and CC-124 were obtained from the Chlamydomonas Resource Center (University of Minnesota, USA) while T222^+^ and the *pgr5* mutant strain were kindly provided by Prof. Michael Hippler (University of Münster, Germany). All strains were initially grown in Tris-acetate-phosphate (TAP) medium for 3 days at 23 °C under continuous shaking and constant illumination at an intensity of 80–100 µmol photons m^−2^ s^−1^ (white fluorescent light). Then, cells were centrifuged at 2900 rpm for 10 min at 20 °C and cells pellets were resuspended in their respective treatment media as described below.

To test the nitrate removal capacity of CC-1690, CC-124, T222^+^ and *pgr5*, cell pellets were resuspended in Tris-phosphate (TP) medium and RAS effluent at an optical density (OD_680_) of 0.05. Cells were then grown for 7 days in 600 ml flasks containing 400 ml of culture at 20 °C under continuous bubbling with room air and illuminated at 100−120 µmol photons m^−2^ s^−1^. Growth curves in each medium, together with nitrate and nitrite removal from RAS effluent, were used to assess the potential of each strain for effluent bioremediation.

To evaluate the H_2_ production capacity of CC-1690 and compare it with that of CC-124, cell pellets were resuspended in high-salt (HS) medium and the Chlorophyll (Chl) content was adjusted to 20 µg Chl (a + b) ml^−1^.

For integrating bioremediation of RAS effluent with H_2_ production, CC-1690 cell pellets were resuspended in RAS effluent and Chl content adjusted to 1 µg Chl (a + b) ml^−1^. Cells were then grown in a Multi-Cultivator MC 1000-OD system (Photon Systems Instruments, Czech Republic) for approx. 70 h at 23 °C under continuous bubbling with CO_2_-enriched air (0.5% v/v) and illuminated at 90 µmol photons m^−2^ s^−1^. Growth rate was calculated from the change in OD_680_ during the exponential growth using the following equation:

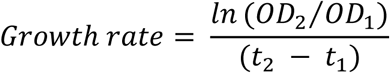

where OD_2_ and OD_1_ represent the OD_680_ at the end (t_2_) and beginning (t_1_) of the cultivation period, respectively (Andersen, 2005).

To assess the H_2_ production capacity of CC-1690 after cultivation in RAS effluent, cultures were centrifuged at 2900 rpm for 10 min at 20 °C and cell pellets were resuspended in their own supernatant to a final Chl content of 20 µg Chl (a + b) ml^−1^.

A previously established protocol for inducing photoautotrophic, anaerobiosis-driven H_2_ evolution was employed (Nagy *et al*., 2018). Briefly, once Chlorophyll (Chl) content was adjusted to 20 µg Chl (a + b) ml^−1^, 30 ml of culture were transferred to glass serum bottles (52 mm × 95 mm, total volume: 120 ml) and sealed with rubber septa under sterile conditions. Approx. 2 g of an iron-salt-based, non-cytotoxic O_2_ absorbent (O2Zero-50 cc loose; Global Reach Ltd, London, UK) was introduced in the headspace of serum bottles to reach an O_2_ concentration below 0.05% in the headspace. A dark anaerobic incubation was then performed by flushing the headspace with N_2_ gas for 15 min and keeping the cultures in darkness for 3 h with constant shaking (120 rpm). Headspaces were then flushed again with N_2_ gas for 15 min before starting H_2_ production under constant light at either 100 or 350 µmol photons m^−2^ s^−1^. Sampling of cultures took place before the start of the dark anaerobic incubation (control, aerobe samples), after the 3 h dark anaerobic incubation, and after 24, 48 and 72 h of H_2_ production.

### 2.3. H_2_ quantification

The H_2_ concentration of the headspace of serum bottles was determined by collecting a 200 µl aliquot with a gas-tight Hamilton microsyringe. Headspace samples were manually injected into a Hewlett Packard 5890 gas chromatograph equipped with an HP-PLOT Molesieve column (30 m × 0.53 mm × 0.25 µm) maintained at 40 °C and coupled to a thermal conductivity detector operated at 160 °C. Argon served as the carrier gas at a linear velocity of 115 cm s^−1^. Following each 24 h gas sampling interval, the headspace of serum bottles was purged with N_2_ gas to prevent H_2_ accumulation exceeding 5% (Kosourov et al., 2012).

### 2.4. Chl *a* fluorescence measurements

Before the measurements, *C. reinhardtii* cultures were dark-adapted for 10 mi. Subsequently, 60 µl of cell suspension was placed onto a Whatman glass microfibre filter (GF/B) and a HandyPEA instrument (Hansatech Instruments Ltd, UK) was used to record the Chl *a* fluorescence signals. The 24, 48 and 72 h H_2_ production samples were placed under moderate light intensity and ambient atmosphere for 2 h upon opening of serum bottles to allow cells to recover before measurement.

### 2.5. Immunoblot analysis

At each time point, 2 ml of culture was collected, spun down to remove the supernatant, and frozen in liquid nitrogen. Immunoblotting was carried out as described in Vidal-Meireles *et al*. (2022) with minor changes. Briefly, a cell extract equivalent to 0.5 million cells was mixed with 6x Laemmli buffer (375 mM Tris/HCl [pH 6.8], 60% [v/v] glycerin, 12.6% [w/v] sodium dodecyl sulfate, 600 mM dithiothreitol, 0.09% [w/v] bromophenol blue) and incubated at 75 °C for 10 min before loading. Protein separation and western blot were carried out as described in Podmaniczki et al., (2021). Specific polyclonal antibodies (produced in rabbit) against PsbA, PSBO, CP47, PetB and PsaA were purchased from Agrisera AB (Sweden). Densitometry analysis of the immunoblots were carried out using ImageJ (v1.54p).

### 2.6. Pigment analysis

Chl (a + b) content of cultures was determined according to Porra *et al*. (1989). For pigment analysis, alga cells were harvested by centrifugation, and the pellets were frozen in liquid N_2_ and stored at -80 °C until usage. HPLC analysis was carried out as previously described (Nagy *et al*., 2025). Briefly, frozen samples were resuspended in 1 ml of extraction buffer consisting of 80 % acetone and 20 % methanol. Pigments were extracted for 30 min at 25 °C in the dark with continuous shaking at 1000 rpm. The extract was centrifuged at 11500 g for 10 min at 4 °C, the supernatant was collected and filtered through a 0.22 µm PTFE syringe filter. Quantification of carotenoids was performed by HPLC using a Prominence HPLC system (Shimadzu, Kyoto, Japan). Chromatographic separations were performed on an Acclaim reverse-phase C30, 3 µm, 150 x 4.6 mm column (Thermo Fisher Scientific Inc.). Pigments were eluted using a linear gradient from 100% solvent A (acetonitrile:water:trimethylamine 90:10:0.1) to 40% solvent ‘A’ and 60% solvent ‘B’ (ethyl acetate). Pigments were identified based on their retention time and then quantified by the integrated chromatographic peak area recorded at the wavelength of maximum absorbance for each type of pigment.

The de-epoxidation index was calculated as the molar ratio of the de-epoxidated and the total xanthophyll cycle components using the following formula:

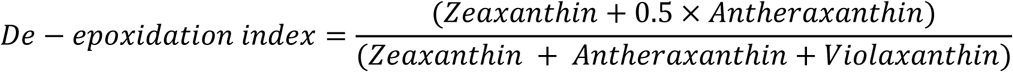

### 2.7. Biomass analysis

At each sampling point, cultures were centrifuged at 15000 rpm at 4 °C for 3 min, the supernatant was discarded, and the samples were flash frozen in liquid N_2_ before being stored at −80 °C. Samples were then freeze-dried overnight before analysis. For total carbon and nitrogen analysis, approx. 1 mg of dry biomass was weighed in a smooth-wall tin cup (D4057 Elemental Microanalysis). Tin cups containing biomass were left in a desiccator until carbon and nitrogen content was analysed using a Thermo Finnigan DELTAplusAdvantage mass spectrometer (Thermo Electron) connected to a FlashEA 1112 Elementar Analyser. Birch leaves (*Betula pendula*) were used as internal standards during the run. Conversion of total nitrogen to protein content was carried out using a conversion factor of 4.08 (Templeton & Laurens, 2015). For total lipid quantification, approx. 2.2 mg of dry biomass were transferred to a 10 ml Kimax tube. Total lipids were then extracted using 3.75 ml chloroform/methanol/water (4:2:1) and sonication (10 min). After phase separation, the lipid-rich lower phase was transferred to a new Kimax tube and solvents were evaporated at room temperature under a N_2_ stream. Dried lipids were resuspended in150 µl of chloroform and transferred to pre-weighed smooth-wall tin cup (D4057 Elemental Microanalysis). Solvents were evaporated at room temperature and total lipid content was calculated as the change in weight in tin cups. For fatty acid analysis approx. 1 mg of dry biomass was used, and total lipids were extracted as described above. After phase separation and evaporation of solvents, lipids were dissolved in 1 ml toluene, and fatty acids were transesterified overnight at 50 °C using 2 ml of methanolic H_2_SO_4_ (1%, v/v). Fatty acid methyl esters were separated by the addition of 2 ml hexane, transferred to a new Kimax tube and hexane was evaporated under a N_2_ stream. Samples were resuspended in 300 µl of hexane and transferred to vials before being analysed with a gas chromatograph equipped with a flame ionization detector (FID-2010 Plus, Shimadzu) using a DB-FastFAME column (30 m × 0.25 mm × 0.25 μm; Agilent) and a previously described temperature program (Calderini *et al*., 2023). Identification of fatty acids was done using retention times of an external fatty acid standard (GLC reference standard 556C, Nu-Chek Prep, Elysian) and quantification was based on peak integration using a three-point calibration curve using the same external standard. Fatty acid losses during the extraction process were accounted for by using the internal standard free FA C23:0 (Larodan) which was added to all samples before lipid extraction.

### 2.8. Nutrient analysis

Water samples were collected, filtered through 0.45 µm pore size nylon syringe filters (Whatman), and stored at −20 °C. Nutrient analysis of samples (nitrate + nitrite, phosphate, and ammonium) was conducted by KVVY Tutkimus Oy (Finland).

### 2.9. Statistical analysis

When applicable, treatment effect was first tested using one-way ANOVA or Kruskal-Wallis test depending on the equality of variances assessed with Levene’s test. If the treatment effect was significant, Dunnett’s or Dunn’s post-hoc tests were used to compare the initial conditions with the rest of the treatments (p < 0.05). The presented data represent at least three independent experiments, and the exact number of biological repetitions are indicated in the figure captions. All statistical analyses were conducted using R language (v4.5.1) with RStudio (v2025.09.2).

## 3. Results and discussion

### 3.1. H_2_ production and nutrient remediation by nitrate-assimilating *C. reinhardtii* CC-1690

For this study, we selected the *C. reinhardtii* CC-1690 strain, which possesses wild-type alleles at the NIT1 and NIT2 loci (Pröschold *et al*., 2005) and can therefore utilize nitrate for growth. To our knowledge, the H_2_ production capacity of this strain has not been previously evaluated; hence we first characterized its photobiological H_2_ evolution potential under conditions comparable to those previously described, namely using anaerobic induction, carbon limitation, and the inclusion of an O_2_ scavenger in the headspace (Nagy *et al*., 2018).

CC-1690 cultures were grown in TAP medium for three days and subsequently transferred to HS medium at a Chl (a+b) concentration of 20 µg ml^−1^. Hydrogenase expression was induced by a 3 h dark anaerobic incubation as described in Nagy *et al*. (2018), after which cultures were exposed to continuous illumination at 100 µmol photons m^−2^ s^−1^. In agreement with previous results (Nagy *et al*., 2018), the inclusion of an O_2_ scavenger in the headspace approx. doubled the H_2_ yield of CC-1690 (Fig. S1). Daily H_2_ production ranged from 13 to 26 µmol H_2_ mg^−1^ Chl, reaching a cumulative total of 61 µmol H_2_ mg^−1^ Chl over 72 h (Fig. S1). When compared with the widely studied CC-124 strain, CC-1690 produced approx. 38% less H_2_ during the 72 h under anaerobiosis-induced, carbon-limited conditions (Fig. 1a,b).

**Fig. 1.**
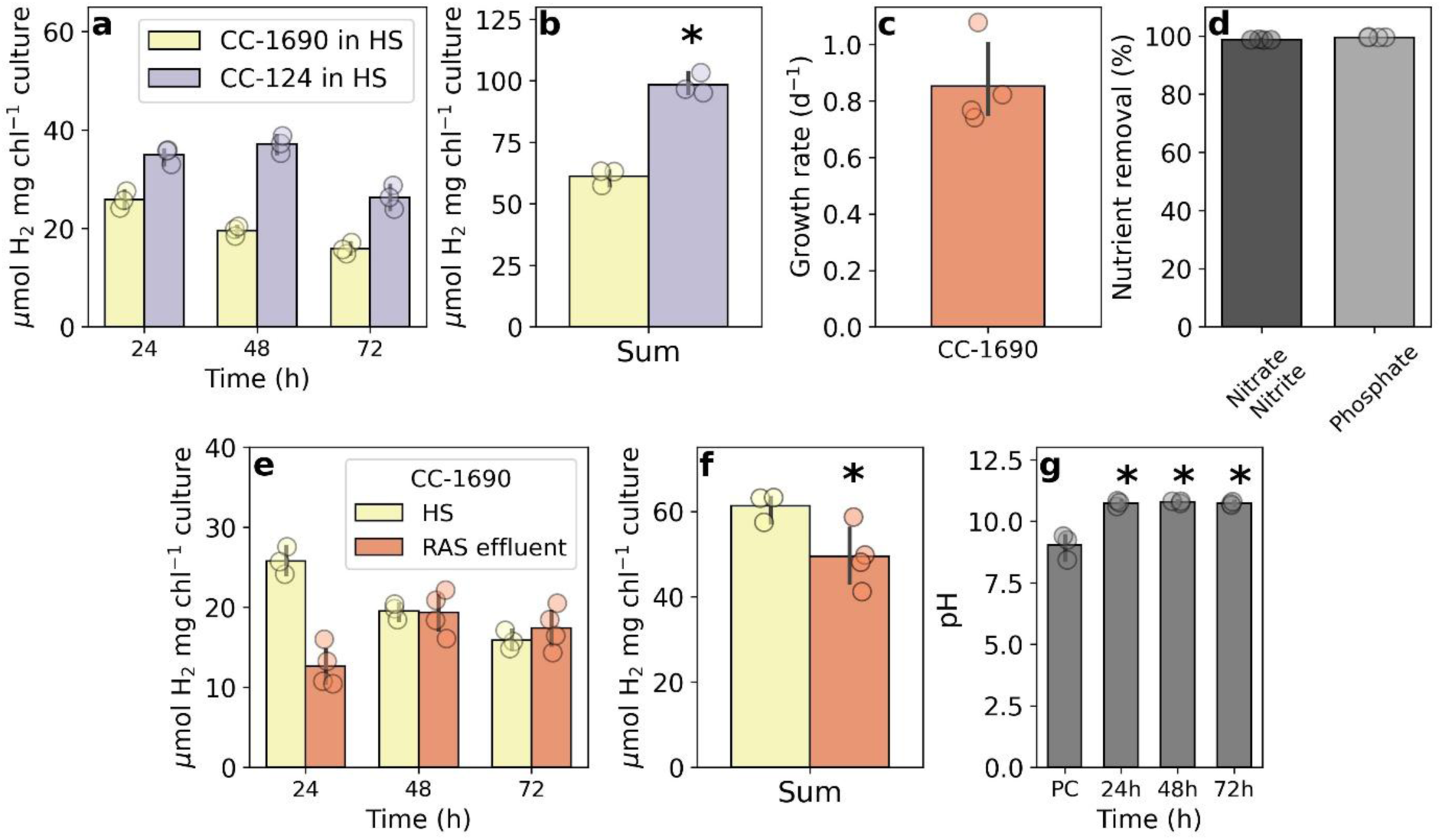
Anaerobiosis-induced H_2_ production by *C. reinhardtii* under continuous light at 100 μmol photons m^−2^s^−1^ and a Chl (a + b) content of 20 µg ml^−1^. Daily (**a**) and cumulative 72 h (**b**) H_2_ production in HS medium for CC-124 and CC-1690. Growth rate (d^−1^) (**c**) and total nutrient (%) removal (**d**) of CC-1690 after approx. 70 h of cultivation in RAS effluent. Daily (**e**) and cumulative 72 h (**f**) H_2_ production in RAS effluent and HS medium for CC-1690. RAS effluent pH (**g**) post-cultivation of CC-1690 (PC) and at 24, 48, and 72 h of H_2_ production. The data represent means and standard deviation of at least three independent biological replicates. Asterisks indicate significant difference in the total H_2_ produced based on one-way ANOVA (p < 0.05).

The bioremediation performance of CC-1690 was then tested by cultivating the strain in RAS effluent. CC-1690 exhibited robust growth (Fig. S2), whereas the best-performing H_2_-producing strains reported to date (CC-124, T222⁺, and *pgr5*; Steinbeck *et al.,* 2015; Nagy *et al.,* 2021) were unable to assimilate nitrate and nitrite from the RAS effluent. Stress-related phenotypes were observed in these strains after a few days of cultivation and only upon the addition of ammonium the cells were able to recover (Fig. S2). Under optimized growth conditions, CC-1690 reached a growth rate of approx. 0.85 day^−1^ and removed 99% of the available nitrate, nitrite, and phosphate from RAS effluent within approx. 70 h of cultivation (Fig. S2; Table 1; Fig. 1c,d). The observed growth rate was comparable to that of the fast-growing *Chlorella vulgaris* (Hiltunen *et al.,* 2025) and higher than other tested species grown in similar RAS effluent (Calderini *et al.,* 2021; Timilsina *et al*., 2025).

**Table 1.** Chemical characteristics of filtered (0.22 µm) UV-sterilized RAS effluent prior and after cultivation of *C. reinhardtii* CC-1690 for approx. 70 h and after 72 h of H_2_ production under continuous light at 100 μmol photons m^−2^s^−1^. Post-cultivation nutrient values represent the average of eight independent biological replicates, while values after 72 h of H_2_ production represent the average of four independent biological replicates.

|  | RAS effluent | Post-cultivation | 72 h H<sub>2</sub> production |
| --- | --- | --- | --- |
| Ammonium (mg l<sup>-1</sup>) | 2.0 | 3.9 | 5.0 |
| Phosphate (mg l<sup>-1</sup>) | 1.1 | 0.005 | 0.018 |
| Nitrate + nitrite (mg l<sup>-1</sup>) | 25 | 0.3 | <0.2 |
| N:P ratio * | 24.5 | 840 | 288.9 |
| pH | 7.31 | 9.02 | 10.72 |

\* Calculated as the ratio of nitrogen to phosphate containing compounds in mg l<sup>-1</sup>.

To avoid transferring cells into fresh HS medium, we tested the capacity of CC-1690 to produce H_2_ directly in RAS effluent following cultivation and dark-anaerobic incubation. Daily H_2_ production ranged from approx. 13 to 18 µmol H_2_ mg^−1^ Chl, reaching a total of approx. 49 µmol H_2_ mg^−1^ Chl after 72 h (Fig. 1e,f). Compared to HS medium, H_2_ production was lower in RAS effluent, primarily due to reduced output during the first 24 h of illumination (Fig. 1e,f). The pH of RAS effluent increased by approx. 3.4 units after 24 h of H_2_ production relative to post-cultivation conditions (Fig. 1g). Notably, the use of higher light intensities (350 μmol photons m^−2^s^−1^) did not increase H_2_ production, yielding approx. 46 µmol H_2_ mg^−1^ Chl after 72 h (Fig. S3).

It is likely that the difference in H_2_ production between RAS effluent and HS medium originated from residual inorganic carbon and nitrate present in the medium. Inorganic carbon availability activates the Calvin-Benson cycle, which diverts electron flux toward carbon fixation at the expense of the hydrogenase (Gaffron & Rubin,1942). Removal of inorganic carbon from the medium due to carbon fixation would partially explain the increase in pH after 24 h of continuous light (Fig. 1g). Additionally, although present at low concentrations (Table 1), nitrate and nitrite suppress H_2_ production and H_2_ evolution can occur only upon their depletion (Aparicio *et al*., 1985).

Potential strategies to improve H_2_ yields in CC-1690 cultivated in RAS effluent include the use of photobioreactors with improved light-surface ratio (Nagy *et al*., 2024). Alternatively, genetic engineering of the CC-1690 strain to suppress competing electron sinks downstream of PSI, including cyclic electron transport and flavodiiron proteins, is a potential avenue to enhance H_2_ production (Steinbeck *et al*., 2015; Jokel *et al*., 2019).

### 3.2. Changes in the photosynthetic apparatus and HydA during the H_2_ production phase

Next, we examined the response of the photosynthetic apparatus to the H_2_ production conditions in RAS effluent. Anaerobiosis-induced H_2_ production led to a small but progressive daily decrease in the Chl content relative to the initial Chl concentration. After 48 h, the Chl content decreased by approx. 10%, with losses reaching 14% after 72 h compared to the initial levels (Fig. 2a). Fast Chl *a* fluorescence (OJIP), a diagnostic tool for detecting alterations in the photosynthetic electron transport chain (Schansker *et al*., 2014), revealed a progressive attenuation in intensity and substantial changes in its kinetics (Fig. 2b). This effect was observed even though the 24, 48 and 72 h samples underwent a 2 h incubation under standard growth conditions (moderate light intensity and ambient atmosphere) prior to measurement, a step designed to enable the recovery of the electron transport chain following reversible downregulation (Nagy *et al*., 2024). The F_V_/F_M_ value, an indicator of PSII efficiency, declined from approx. 0.65 post-cultivation to 0.47 at 48 h, and finally to 0.33 at 72 h (Fig. 2c).

**Fig. 2.**
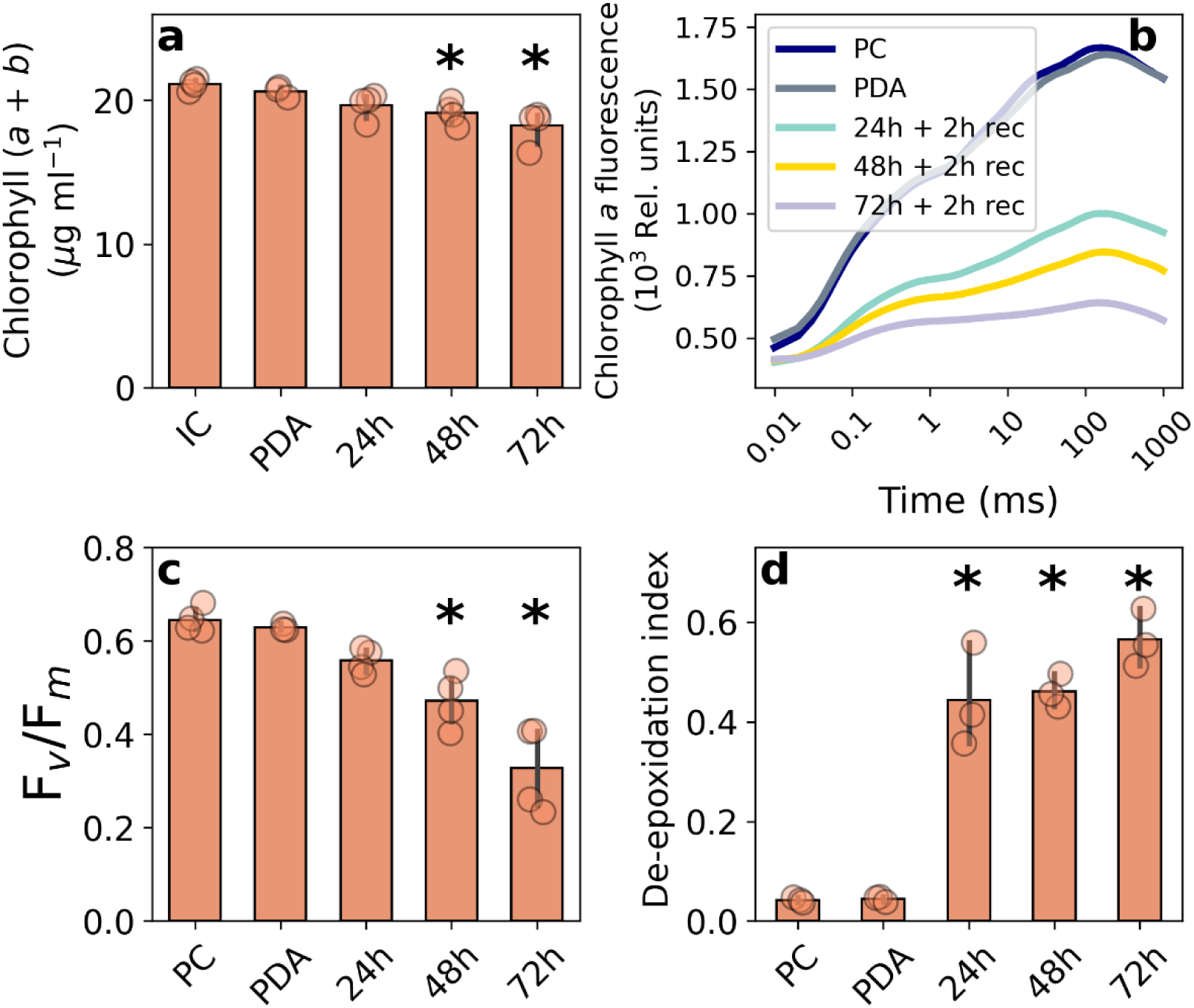
Characterization of the photosynthetic apparatus and pigment composition of CC-1690 before and during anaerobiosis-induced H_2_ production under continuous light at 100 μmol photons m^−2^s^−1^. Chl (a + b) content (**a**), fast Chl *a* fluorescence transient (OJIP) (**b**), F_V_/F_M_ values derived from the OJIP transient (**c**), and de-epoxidation index (**d**) were analysed for the starting conditions, after dark anaerobic incubation (PDA) and 24, 48 and 72 h of H_2_ production. Given that the Chl (a + b) concentration was adjusted to 20 µg ml^−1^ in serum bottles prior to H_2_ production, such Chl content was considered the initial conditions (IC), while all other starting measurements were taken immediately post-cultivation (PC) in RAS effluent. The data represent means and standard deviation of at least three independent biological replicates. Asterisks indicate a significant difference between the treatment group and the starting conditions (PC or IC) by either Dunnett’s or Dunn’s post-hoc test following a significant one-way ANOVA or Kruskal-Wallis test (p < 0.05).

Given that light stress may lead to the conversion of violaxanthin to zeaxanthin in *C. reinhardtii* (Erickson *et al*., 2015), we calculated the degree of violaxanthin cycle de-epoxidation during the H_2_ production phase. The de-epoxidation index increased 10-fold after 24 h of light exposure and reached its maximum after 72 h, exhibiting an approx. 13-fold increase relative to post-cultivation and post-dark incubation levels (Fig. 2d). Significant increases in the Chl a/b ratio and the lutein content (normalized to Chl *a*) after light exposure further suggest the occurrence of light stress during the H_2_ production phase (Fig. S4). Additionally, the increase in lutein coincides with a decrease in loroxanthin (Fig. S4), suggesting the activation of the loroxanthin cycle (Van Den Berg and Croce, 2022). These changes in pigment composition are consistent with a reduction in light-harvesting antenna size and an increase in non-photochemical quenching (Dupuis *et al*., 2025).

To investigate whether key components of the electron transport chain were affected by the anaerobiosis-induced H_2_ production, immunoblot analysis was carried out. The relative amounts of PSII subunits responded differently to the experimental conditions: PsbA levels were maintained, whereas the CP47 content increased by approx. 40% and PSBO decreased by approx. 60% after 72 h compared to the initial conditions (Fig. 3a–c). The cytochrome b_6_f subunit PetB was maintained, while the PSI subunit PsaA exhibited some variation (Fig. 3e). Collectively, these results confirm that light stress occurred in CC-1690 under the applied conditions. This conclusion is further supported by the observation that higher light intensities (350 μmol photons m^−2^ s^−1^) during H_2_ production caused more pronounced reductions in Chl content and F_V_/F_M_ values after 72 h, accompanied by corresponding changes in pigment profiles and protein contents (Fig. S5, Fig. S6).

**Fig. 3.**
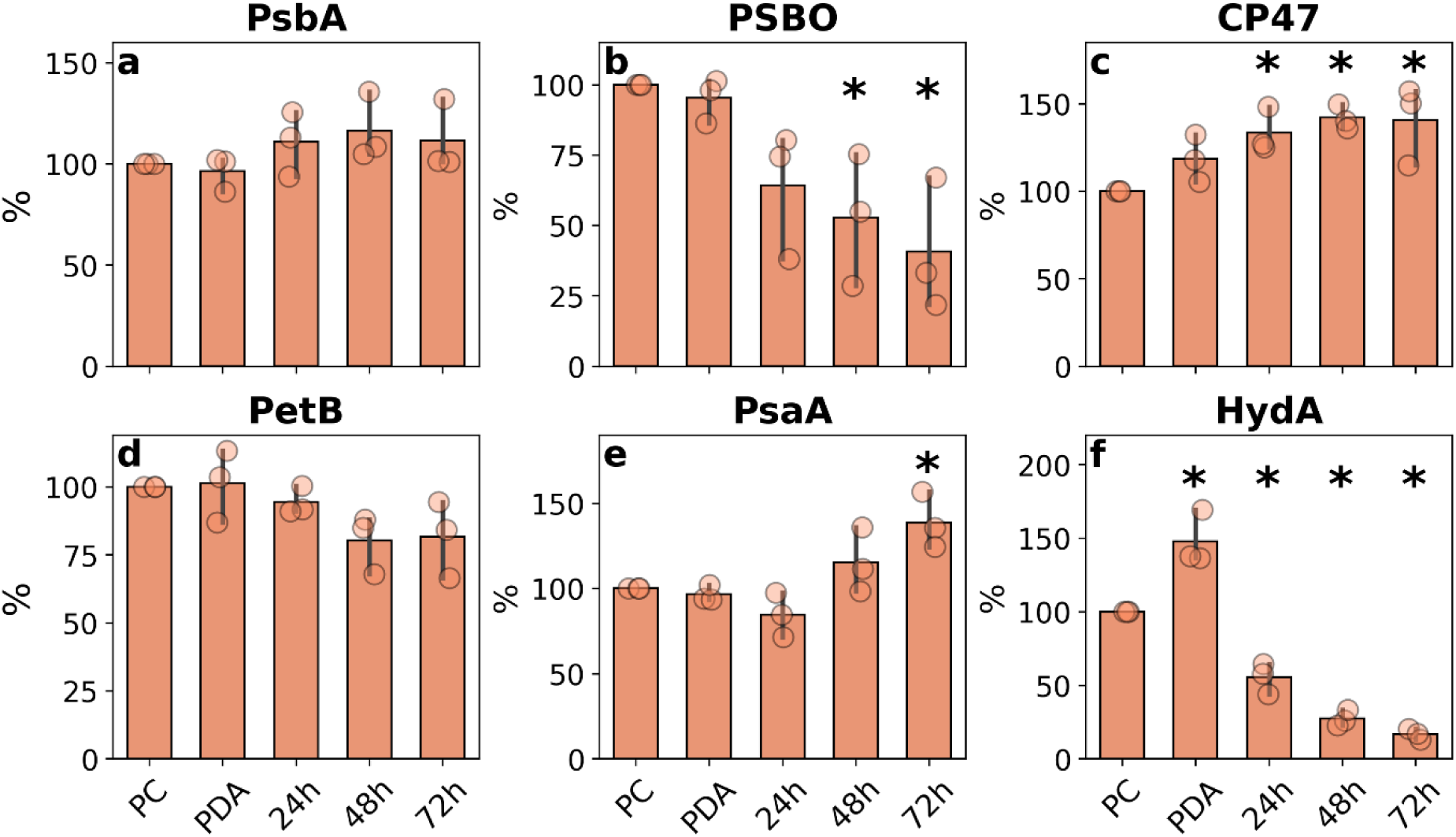
Immunoblot analysis for the semi-quantitative determination of key components of the electron transport chain and HydA of CC-1690 before and during anaerobiosis-induced H_2_ production under continuous light at 100 μmol photons m^−2^s^−1^. The relative abundance of the PSII subunits PsbA (**a**), PSBO (**b**), CP47 (**c**); the cytochrome b_6_f PetB (**d**); the PSI subunit PsaA (**e**); and the hydrogenase HydA (f) were analysed. Samples were taken post-cultivation (PC) in RAS effluent, post-dark anaerobic incubation (PDA) and after 24, 48 and 72 h of H_2_ production. The data represent means and standard deviation of three independent biological replicates. Data are expressed as percentage (%) normalized to PC conditions. Asterisks indicate a significant difference between the treatment group and the PC conditions by either Dunnett’s or Dunn’s post-hoc test following a significant one-way ANOVA or Kruskal-Wallis test (p < 0.05).

The amount of hydrogenase HydA was maximal after the dark anaerobic incubation, but decreased sharply upon light exposure to levels lower than those observed post-cultivation, retaining only approx. 17% of the initial level after 72 h (Fig. 3f). Such a reduction of HydA has been previously observed under similar conditions in the CC-124 strain (Nagy *et al*., 2021, Nagy *et al*., 2024), and is most likely due to the inhibitory effect of the small amounts of O_2_ inevitably generated in the chloroplasts by PSII activity.

### 3.3. Effects of H_2_ production on CC-1690 biomass quality

Recovery of high-quality microalgal biomass following the H_2_ production phase is critical for the integration of downstream applications. Traditional methods, such as the use of sulphur deficiency to promote H_2_ evolution, lead to a stress-induced metabolic shift that decreases the quality of the resulting biomass by lowering protein content (Kosourov *et al*., 2002, Matthew *et al*., 2009). Hence, we investigated whether CC-1690 biomass is altered during and after the H_2_ production phase. On a dry weight basis, the post-cultivation biomass was composed of approx. 50% carbon, 25% protein, and 21% lipids. (Fig. 4a-c). Total carbon and protein content were not significantly affected by the dark anaerobic incubation or by the H_2_ production conditions.

**Fig. 4.**
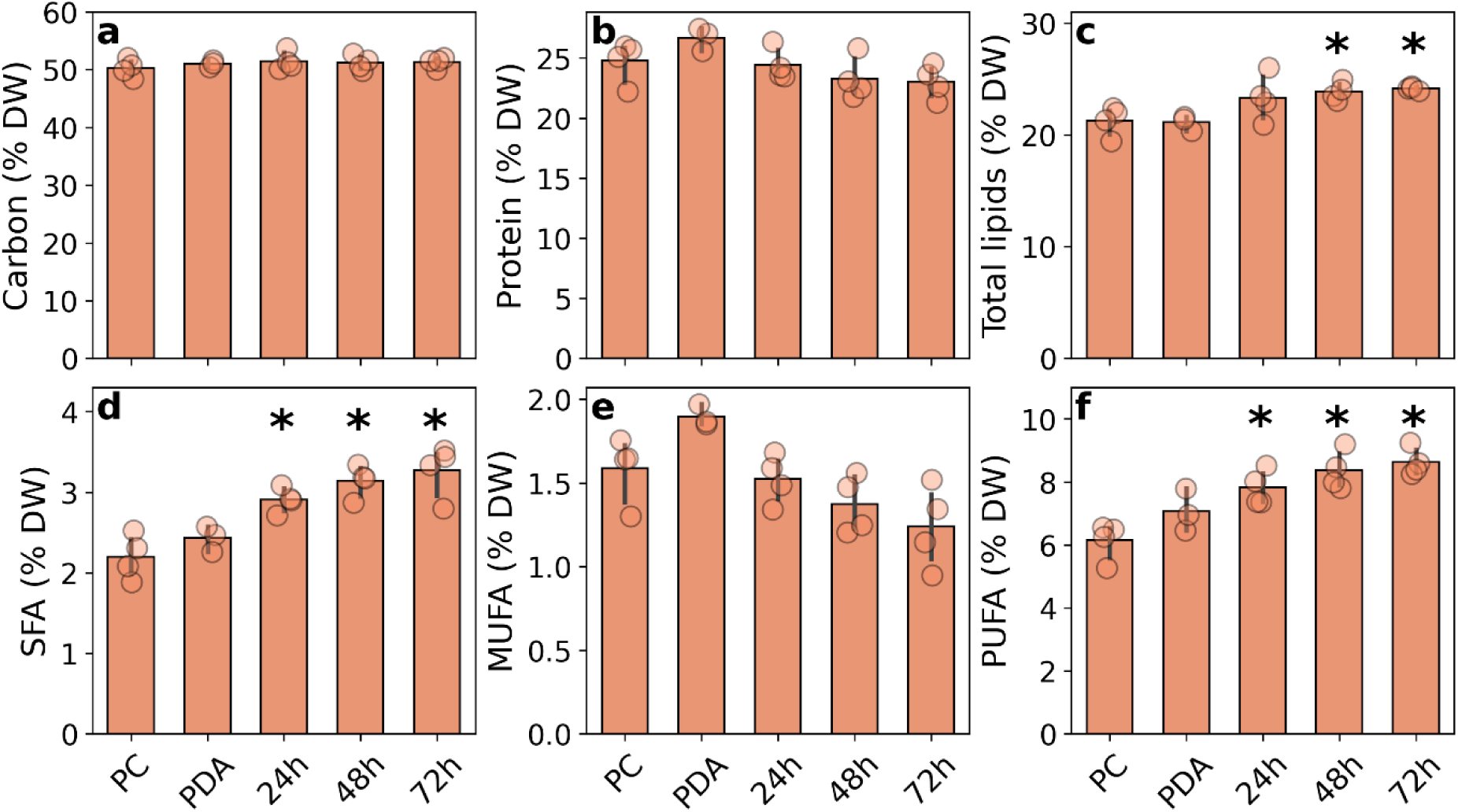
Biomass composition of CC-1690 before and during anaerobiosis-induced H_2_ production under continuous light at 100 μmol photons m^−2^s^−1^. Total carbon (**a**), protein (**b**), and lipid (**c**) contents, as well as saturated (SFA) (**d**), monounsaturated (MUFA) (**e**) and polyunsaturated (PUFA) (**f**) fatty acids, expressed as a percentage of dry weight (DW). Samples were taken post-cultivation (PC) in RAS effluent, post-dark anaerobic incubation (PDA), and after 24, 48 and 72 h of H_2_ production. The data represent means and standard deviation of three independent biological replicates. Asterisks indicate a significant difference between the treatment group and the PC conditions by either Dunnett’s or Dunn’s post-hoc test following a significant one-way ANOVA or Kruskal-Wallis test (p < 0.05).

Total lipid content increased slightly (approx. 3%) upon 48 and 72 h of illumination (Fig. 4c), driven by the accumulation of fatty acids (Table S1; Fig. 4d-f). Saturated (SFA) and polyunsaturated fatty acids (PUFA) increased by approx. 49% and 41%, respectively, after 72 h of H_2_ production compared to the post-cultivation conditions (Fig. 4c,f). Monounsaturated fatty acids (MUFA) remained largely unchanged during the H_2_ production phase. The changes in SFA were predominantly a result of increases in palmitic acid (16:0), which accounted for approx. 20% of all fatty acids, while the increase in PUFA was primarily driven by α-linolenic acid (18:3ω-3), which accounted for approx. 26% of all fatty acids (Fig. S7).

Due to the complexity of the conditions to which CC-1690 was exposed to during the H_2_ production phase, it is difficult to link the observed changes in biomass to specific metabolic pathways. During the H_2_ production phase, the nitrogen concentration in RAS effluent was negligible, suggesting a possible metabolic response to nitrogen deprivation by CC-1690. Such response usually leads to the degradation of Chl and protein and the initial channelling of CO_2_ fixation towards lipid production (Saroussi *et al*., 2016; Arora *et al*., 2018). However, due to the substrate limitation of the Calvin-Benson cycle and the expression of HydA, only the degradation of macromolecules could provide carbon for lipid synthesis under the tested conditions. Although we did not observe gross alterations in the protein content typical of nitrogen deprivation studies in *C. reinhardtii* (Schmollinger *et al.,* 2014; Arora *et al*., 2018), it is likely that the degradation of a small fraction of proteins and other macromolecules were responsible for fuelling the observed increase in lipid content.

Altogether, the biomass analysis supports the potential valorisation of this biomass as a biofuel feedstock due to the high content of palmitic acid, while the stable protein content and elevated proportion of α-linolenic acid highlight the suitability of the biomass for aquafeed applications (Senadheera *et al*., 2011).

## 4. Conclusion

Bioremediation of RAS effluents using the nitrate-assimilating *C. reinhardtii* strain CC-1690 can be effectively coupled with H_2_ production for further valorisation of the process. Within 70 h of cultivation, CC-1690 achieved near-complete removal of all nitrate and phosphate from the RAS effluent. Post-cultivation induction of H_2_ evolution through anaerobiosis-induced photoautotrophic conditions sustained H_2_ production for 72 h, yielding 49 µmol H_2_ mg⁻¹ Chl. Although indicators of light stress were observed, major photosynthetic subunits remained largely intact, and biomass quality was preserved. These findings highlight the potential of integrating H_2_ production into microalgal treatment of RAS effluents as a viable strategy for simultaneous nutrient removal, bioenergy production, and biomass valorisation.

## Data statement

All data required to replicate the results presented in this study will be available at a provided DOI upon publication. A comprehensive description of the content of each file can be found in the same DOI under the name “Metadata file.”

## CRediT authorship contribution statement

**Marco L. Calderini**: Writing – original draft, Methodology, Formal analysis, Conceptualization. **Valéria Nagy**: Writing – review & editing, Methodology, Formal analysis. **Szilvia Z. Tóth**: Writing – review & editing, Methodology, Supervision, Resources. **László Kovács**: Writing – review & editing, Methodology. **Soujanya Kuntam**: Writing – review & editing, Methodology. **Pauliina Salmi**: Writing – review & editing, Supervision, Project administration, Methodology, Conceptualization.

## Acknowledgements

The authors would like to thank laboratory technicians Éva Herman and Emma Pajunen for their help during the experimental work. This work was funded by the European Union – NextGenerationEU and the Research Council of Finland research grant awarded to P.S. (grant no. 352764) and by the Lendület Program of the Hungarian Academy of Sciences (LP2024/21) awarded to S.Z.T.

