## Supplemental material for "Coupling microalgal bioremediation of recirculating aquaculture effluents with photobiological hydrogen production"

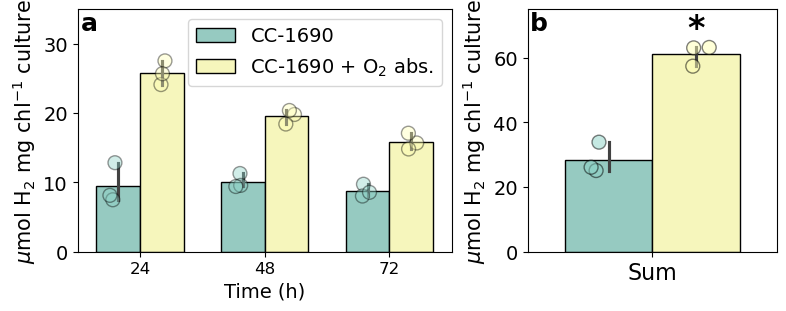

**Figure S1.** Daily (**a**) and 72 h total (**b**) anaerobiosis-induced, carbon-limited H_2_ production by *C. reinhardtii* CC-1690 with or without the addition of an iron-salt-based O_2_ absorbent under continuous light at 100 μmol photons m^−2^s^−1^ and a Chl (a + b) concentration of 20 µg ml^−1^. The data represent means and standard deviation of three independent biological replicates. The asterisk indicates significant difference in the 72 h total H_2_ produced based on one-way ANOVA (p < 0.05).

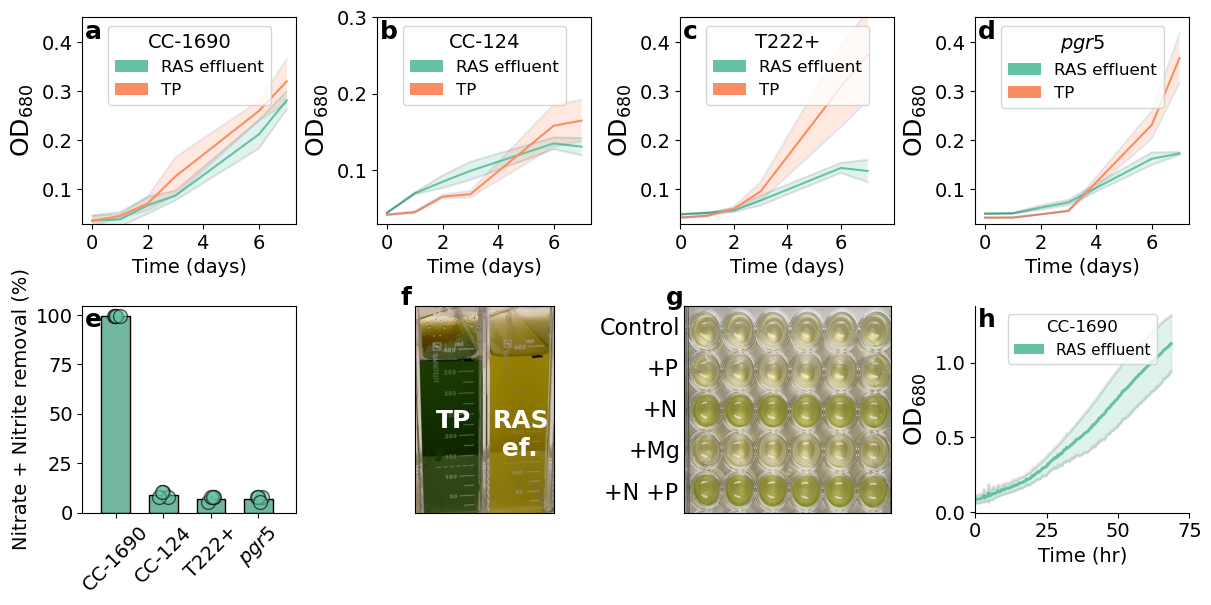
**Figure S2.** Growth and nutrient removal of *C. reinhardtii* wild-type strains CC-1690, CC-124 and T222+, together with *pgr5* mutant, when cultured in RAS effluent and Tris-phosphate (TP) medium. Optical density (OD_680_) (**a**-**d**) measured during a 7-day cultivation period to test the nitrate and nitrite removal capacity of strains. Nitrate and nitrite removal (%) from RAS effluent (**e**). Visual difference in culture color (**f**) of T222+ after 7 days of cultivation in TP medium and RAS effluent (RAS ef.). Post-cultivation cultures of CC-124, T222+ and *pgr5* presented similar coloration difference between cultivation in TP and RAS effluent. Recovery test (**g**) to identify if the addition of phosphate (P), ammonium (N), or magnesium (Mg) to T222+ cells post-cultivation in RAS effluent restores culture coloration. Columns in the well plate represent technical replicates of the same treatment. CC-124, T222+ and *pgr5* identical responses to N addition. Growth curve (OD_680_) of *C. reinhardtii* CC-1690 (**h**) when cultivated in RAS effluent at 23 °C under continuous bubbling with CO_2_-enriched air (0.5% v/v) and illuminated at 90 µmol photons m^−2^ s^−1^. The data represent means and standard deviation of either four (**a**-**e**) or eight (**h**) independent biological replicates.

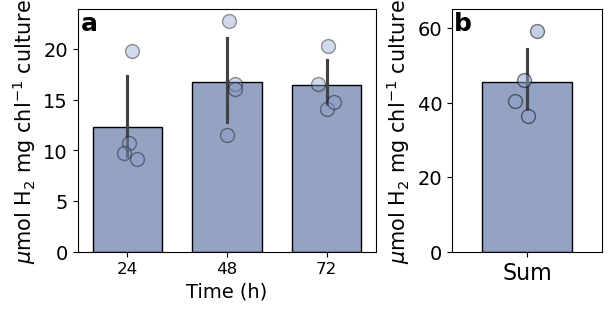

**Figure S3.** Anaerobiosis-induced H_2_ production by *C. reinhardtii* CC-1690 under continuous light at 350 μmol photons m^−2^s^−1^ and a Chl (a + b) concentration of 20 µg ml^−1^. Daily (**a**) and 72 h cumulative (**b**) H_2_ production in RAS effluent after cultivation in the same medium. The data represent means and standard deviation of four independent biological replicates.

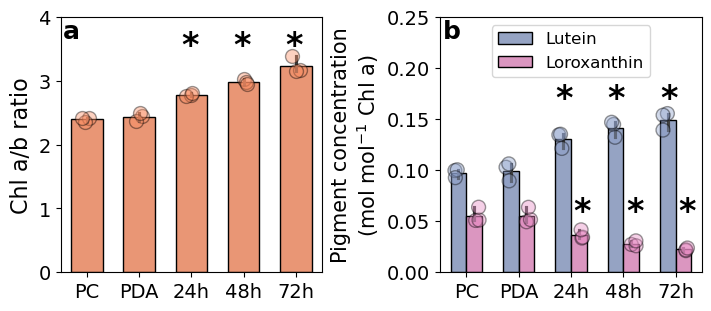

**Figure S4.** Chl a/b ratio (**a**) and lutein content per mol of Chl *a* (**b**) of *C. reinhardtii* CC-1690 before and during anaerobiosis-induced H_2_ production under continuous light at 100 μmol photons m^−2^s^−1^ and a Chl (a + b) concentration of 20 µg ml^−1^. The data represent means and standard deviation of three independent biological replicates. Asterisks indicate a significant difference between the treatment group and the starting conditions (PC or IC) for each pigment by Dunnett's post-hoc test following a significant one-way ANOVA (p < 0.05).

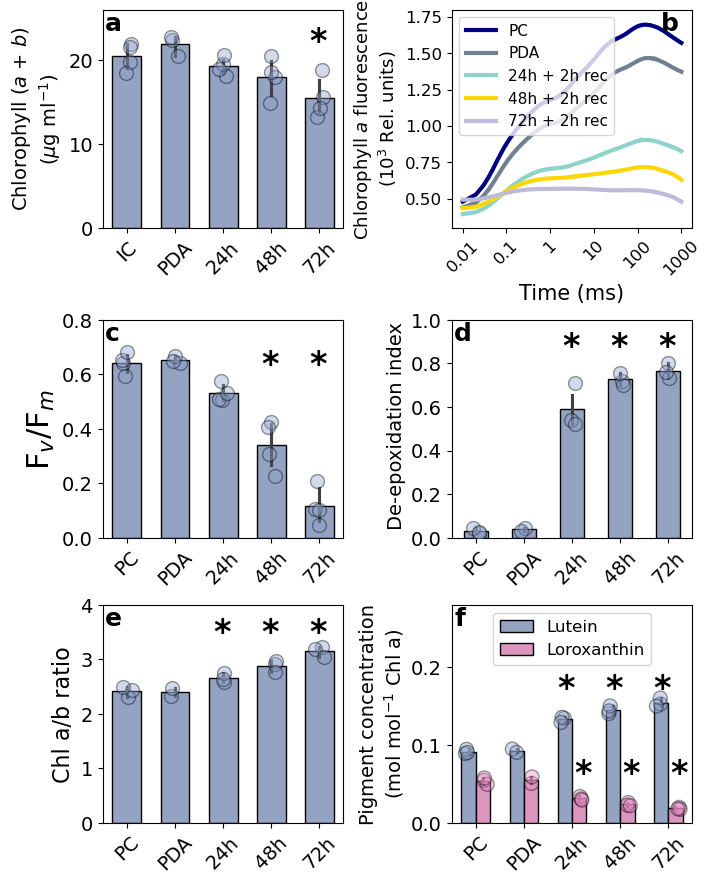

**Figure S5.** Characterization of the photosynthetic apparatus and pigment composition of CC-1690 before and during anaerobiosis-induced H_2_ production under continuous light at 350 μmol photons m^−2^s^−1^ and a Chl (a + b) concentration of 20 µg ml^−1^. Chl (a + b) content (**a**), Chl *a* fluorescence transients (OJIP) (**b**), F_v_/F_m_ values (**c**). De-epoxidation index (**d**), Chl a/b ratio (**e**), and lutein and loroxanthin contents per mol of Chl a (**f**) were analysed for the starting conditions, post-dark anaerobic incubation (PDA), and after 24, 48 and 72 h of H_2_ production. Given that the Chl (a + b) concentration was adjusted to 20 µg ml^−1^ in serum bottles prior to H_2_ production, this value represent the initial conditions (IC), while all other starting measurements were taken immediately post-cultivation (PC) in RAS effluent. The 24, 48 and 72 h samples were placed under moderate light intensity and ambient atmosphere for 2 h upon opening of serum bottles to allow cells to recover before Chl *a* fluorescence measurement. The data represent means and standard deviation of at least three independent biological replicates. Asterisks indicate a significant difference between the treatment group and the starting conditions (PC or IC) by either Dunnett's or Dunn’s post-hoc test following a significant one-way ANOVA or Kruskal-Wallis test (p < 0.05).

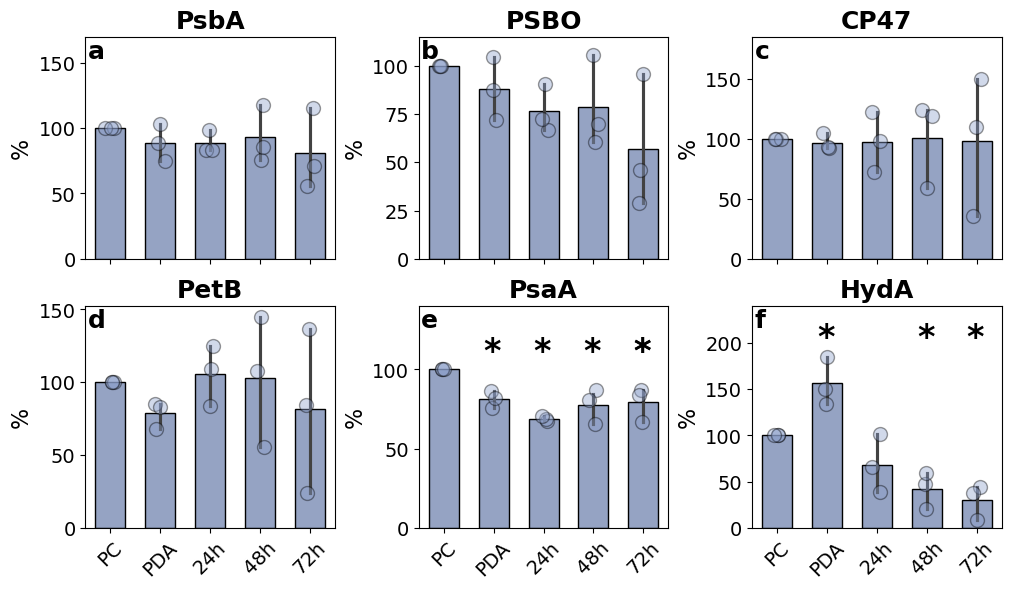
**Figure S6.** Immunoblot analysis for the semi-quantitative determination of key components of the electron transport chain and HydA of CC-1690 before and during anaerobiosis-induced H_2_ production under continuous light at 350 μmol photons m^−2^s^−1^. The relative abundance of the PSII subunits PsbA (**a**), PSBO (**b**), CP47 (**c**); the cytochrome b_6_f subunit PetB (**d**); the PSI subunit PsaA (**e**); and the hydrogenase HydA (f) were analysed. Samples were taken post-cultivation (PC) in RAS effluent, post-dark anaerobic incubation (PDA) and after 24, 48 and 72 h of H_2_ production. The data represent means and standard deviation of three independent biological replicates. Asterisks indicate a significant difference between the treatment group and the PC conditions by either Dunnett's or Dunn’s post-hoc test following a significant one-way ANOVA or Kruskal-Wallis test (p < 0.05).

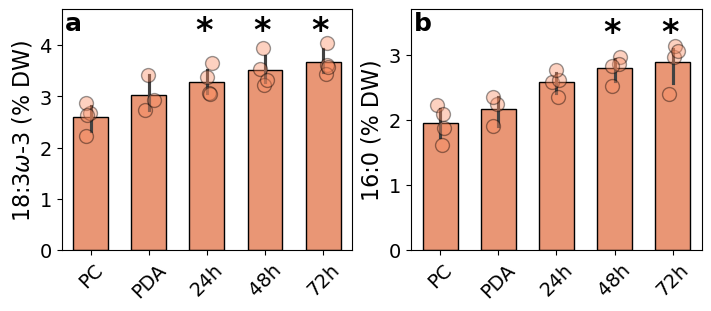

**Fig S7** Percentage per dry weight (DW) of fatty acids α-linolenic acid (18:3ω-3) (**a**) and palmitic acid (16:0) (**b**) before and during anaerobiosis-induced H_2_ production under continuous light at 100 μmol photons m^−2^s^−1^. Samples were taken post-cultivation (PC) in RAS effluent, post-dark anaerobic incubation (PDA) and after 24, 48 and 72 h of H_2_ production. The data represent means and standard deviation of three independent biological replicates. Asterisks indicate a significant difference between the treatment group and the PC conditions by either Dunnett's or Dunn’s post-hoc test following a significant one-way ANOVA or Kruskal test (p < 0.05).

**Table S1** Fatty acid composition of *C. reinhardtii* CC-1690 post-cultivation in RAS effluent (PC), post-dark anaerobic incubation (PDA) and after 24, 48 and 72 h H_2_ production at 100 μmol photons m^−2^s^−1^. Data is expressed as a percentage of dry weight. The data represent means and standard deviation of three independent biological replicates.

|  | **PC** | **PDA** | **24h** | **48h** | **72h** |
| --- | --- | --- | --- | --- | --- |
| 14:0 | 0.02 | 0.03 | 0.03 | 0.04 | 0.04 |
| 16:0 | 1.95 | 2.17 | 2.58 | 2.79 | 2.89 |
| 16:1ω9 | 0.18 | 0.23 | 0.16 | 0.14 | 0.09 |
| 16:1ω7 | 0.16 | 0.2 | 0.15 | 0.16 | 0.16 |
| 16:2ω6 | 0.08 | 0.09 | 0.12 | 0.15 | 0.17 |
| 16:3ω6 | 0.07 | 0.08 | 0.08 | 0.08 | 0.05 |
| 16:3ω3 | 0.25 | 0.28 | 0.36 | 0.38 | 0.39 |
| 16:4ω3 | 1.73 | 2.05 | 2.04 | 2.07 | 2.07 |
| 17:0 | 0.02 | 0.02 | 0.02 | 0.03 | 0.03 |
| 18:0 | 0.19 | 0.2 | 0.25 | 0.27 | 0.29 |
| 18:1ω9 | 1.01 | 1.18 | 0.95 | 0.81 | 0.71 |
| 18:1ω7 | 0.25 | 0.29 | 0.27 | 0.27 | 0.27 |
| 18:2ω6 | 0.65 | 0.67 | 0.92 | 1.04 | 1.08 |
| 18:3ω6 | 0.54 | 0.6 | 0.72 | 0.81 | 0.86 |
| 18:3ω3 | 2.6 | 3.03 | 3.29 | 3.51 | 3.67 |
| 18:4ω3 | 0.23 | 0.27 | 0.3 | 0.33 | 0.34 |
| 20:0 | 0.02 | 0.03 | 0.02 | 0.02 | 0.02 |
| SUM | 9.93 | 11.41 | 12.26 | 12.89 | 13.14 |
| PUFA | 6.15 | 7.07 | 7.83 | 8.37 | 8.63 |
| SFA | 2.2 | 2.44 | 2.91 | 3.14 | 3.27 |
| MUFA | 1.59 | 1.9 | 1.53 | 1.37 | 1.24 |
